# Plant aquaporins: the origin of NIPs

**DOI:** 10.1101/351064

**Authors:** Adrianus C. Borstlap

## Abstract

Many of the aquaporin genes in Cyanobacteria belong to the AqpN-clade. This clade was also the cradle of plant NIPs (nodulin-26 like intrinsic proteins) whose members are transporters for glycerol and several hydroxylated metalloids. The superphylum of Archaeplastida acquired the primordial NIP-gene most likely from the cyanobacterium that, some 1500 million years ago, became the ancestor of all plastids.

---

Nodulin-26 is the major protein in the peribacteroid membrane of soybean nodules. Its coding gene was identified in 1987 and appeared to be related to the gene of the major intrinsic protein of the bovine eye lens and that of the glycerol facilitator of *Escherichia coli* [1,2]. After the protein CHIP28 from erythrocytes joined the club [3] and was characterized as the first water channel or aquaporin protein [4], the family of ‘Major Intrinsic Proteins (MIPs)’ or ‘Aquaporins’ came into view. The protein family consists of two major clades: the clade of aquaporins *sensu stricto*, which function mainly as water channels, and that of the glycerol facilitators (GlpF clade or GIPs, GlpF-like intrinsic proteins). Representatives of both clades are widely distributed in all life forms [5–9].

Plant aquaporins fall into four major subfamilies: plasma membrane intrinsic proteins (PIPs), tonoplast intrinsic proteins (TIPs), nodulin 26-like intrinsic proteins (NIPs), and small, basic intrinsic proteins (SIPs) [9,11]. Two minor groups are the XIPs (X intrinsic proteins), with members in fungi, slime molds and several dicot plant families, and the HIPs (hybrid intrinsic proteins), a small group only present in mosses and lycopods [9,12,13]. GIPs have been encountered in some green algae [14] and in the moss *Physcomitrella patens* [15] but not in vascular plants.

The NIPs are specific for plants. The subfamily is widely represented in the plant kingdom. The number of NIPs range from five in *Physcomitrella* [12] and four in a tiny angiosperm, the duckweed *Spirodela polyrhiza* [16], up to 13 in soybean *(Glycine max)* [17] and 32 in the mesohexaploid oilseed rape *(Brassica napus)* [18]. The absence of NIPs in green algae is remarkable [14].

NIPs are primarily known as transporters for glycerol (C_3_H_5_(OH)_3_) [19] but during plant evolution the proliferation and diversification of NIP genes has resulted in specific transporters for a range of substrates, hydroxylated metalloids in particular [20–22]. NIPs are known for the transport of boron (B(OH)3), silicon (Si(OH)4), arsenic (As(OH)3), and antimony (Sb(OH)3). In addition particular NIPs are transporters for lactic acid (COOH.CH(OH).CH_3_) [23] and aluminium malate, ((Al^3+^)_2_(COO.CH(OH).CH_2_.COO)_3_) [24]. Most telling, perhaps, is the number of 11 NIP-genes in the silicon-accumulating horsetail *Equisetum arvensis*, that are all assumed to code for transporters of silicic acid [25,26].

Although many of the phylogenetic relationships between aquaporins have been elucidated in recent analyses [8, 27–31], the evolutionary origin of the NIPs has remained misty. In reviews it was described as ‘unclear’, ‘unresolved’, ‘uncertain’, or ‘still questioned’ [9,10,22,28]. A first hint for its origin, however, came already in 1996 after the first bacterial water channel protein, AqpZ, was identified [32]. A phylogenetic analysis revealed that NIPs are more similar to the bacterial aquaporin AqpZ than to other plant aquaporins [33]. This has been confirmed in virtually all phylogenetic analyses that followed [8,19,27–31] and has been taken as evidence for a horizontal gene transfer of the ancestral NIP-gene from a bacterium to an ancestor of plants, where it would have replaced a glycerol-transporting GIP [19].

Bacterial aquaporins even more similar to NIPs than the AqpZ-like proteins were first noticed in the group of Urban Johanson [28], and a cyanobacterial aquaporin belonging to a small sister clade of plant NIPs first popped up, just visible, in the extensive review by Federico Abascal *et al.* [see Fig. 1 in ref. 29]. Then an important insight into the diversification of prokaryotic aquaporins was obtained by a thorough phylogenetic analysis of bacterial and archaeal aquaporin sequences by Roderick Nigel Finn and colleagues, which revealed that prokaryote aquaporin genes fall into four clades, the GlpF-, AqpM-, AqpZ- and the AqpN clade [8]. The four clades are represented in both Eubacteria and Archaea [see Fig. S5 in ref. 8]. Bacterial NIP-likes belong to the AqpN clade of prokaryote aquaporins and are the sister group of plant NIPs [28,30]. Since the highest similarities were seen between plant NIPs and AqpN-members of Chloroflexi it has been suggested that the primordial gene of plant NIPs originated from this bacterial phylum [30].

**Figure 1:**
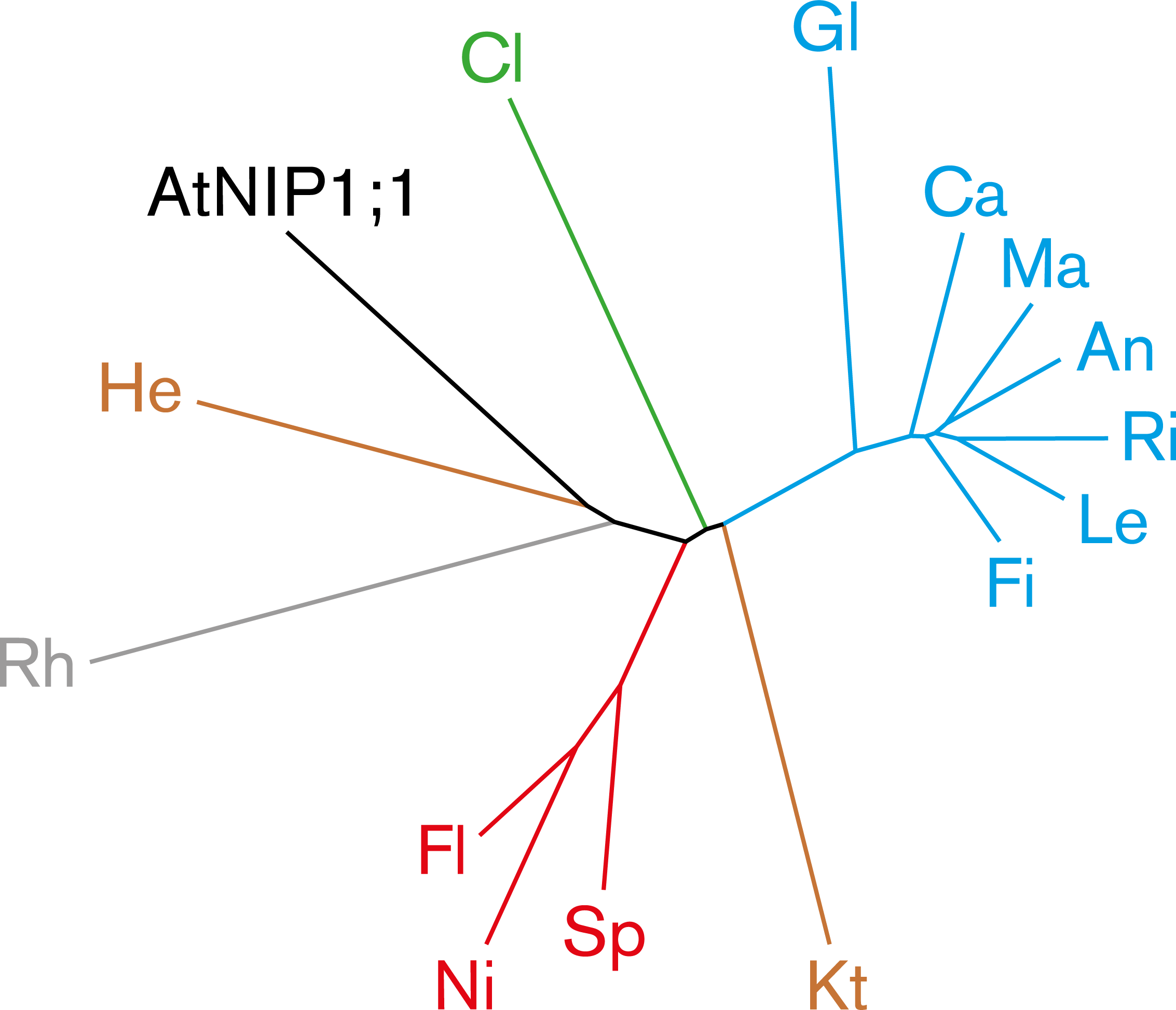
Phylogenetic tree of AtNIP1;1 and its prokaryotic relatives. AtNIP1;1 is one of the nine NIPs from *Arabidopsis thaliana* [11] and belongs, like nodulin-26 of soybean, to the NIP1 branch [41]. The tree shows the most similar proteins from the phylum Cyanobacteria (Gl, *Gloeobacter;* Ca, *Calothrix;* Ma, *Mastigocoleus;* An, *Anabaena;* Ri, *Rivularia;* Le, *Leptolyngbya;* Fi, *Fischerella)* and from all other bacterial phyla (Cl, *Clostridium;* He, *Herpetosiphon;* Rh, *Rhodopirellula;* Fl, *Flavisolibacter;* Sp, *Sphingobacteriales;* Ni, *Niastella;* Kt, *Ktedonobacter).* Color code for the bacterial phyla: blue, Cyanobacteria; green, Firmicutes; brown, Chloroflexi; grey, Planctomycetes; red, Bacteroidetes. Full names of the bacterial species and identification numbers of the proteins can be found in Supplemental Information.

In the AqpN clade shown by Finn and co-workers [Fig. S5 in ref. 8] no cyanobacterial proteins were included, though in the very first publication of a cyanobacterial aquaporin its close relationship to nodulin-26 had already been mentioned explicitly [34]. At present ~300 cyanobacterial aquaporin genes from ~220 cyanobacterial species / strains have been sequenced. Most belong to the AqpZ- or AqpN-clade, a minor part to the GlpF-clade. The origin of the AqpN-clade probably predates the emergence of Cyanobacteria since its most basal species, *Gloeobacter violaceus*, possesses one AqpZ-like and two NIP-like genes.

It is true that NIP-likes from some prokaryotes are more similar to NIPs than their cyanobacterial counterparts. But the difference is rather small and amounts to a few percent points in identical amino acid residues (Table 1). Remarkably, the most similar prokaryotic proteins of a plant NIP fall into various phyla. For example, the seven top hits of a BLASTP search with a NIP from *Arabidopsis thaliana* (AtNIP1;1) as query were from four different phyla: Chloroflexi, Bacteroidetes, Planctomycetes, and Verrucomicrobia (Fig. 1). This may indicate horizontal gene transfer of NIP-like genes between bacteria from various phyla by which it will be very difficult to trace the ancestry of a NIP gene and even wrong inferences about its origin may be made. A striking, hypothetical example is given in Fig. 5 of ref. [35] where it is shown how a plant gene of cyanobacterial descent may be considered, erroneously, to come from a quite unrelated bacterium. Could it be that NIPs have a cyanobacterial origin?

**Table 1:**
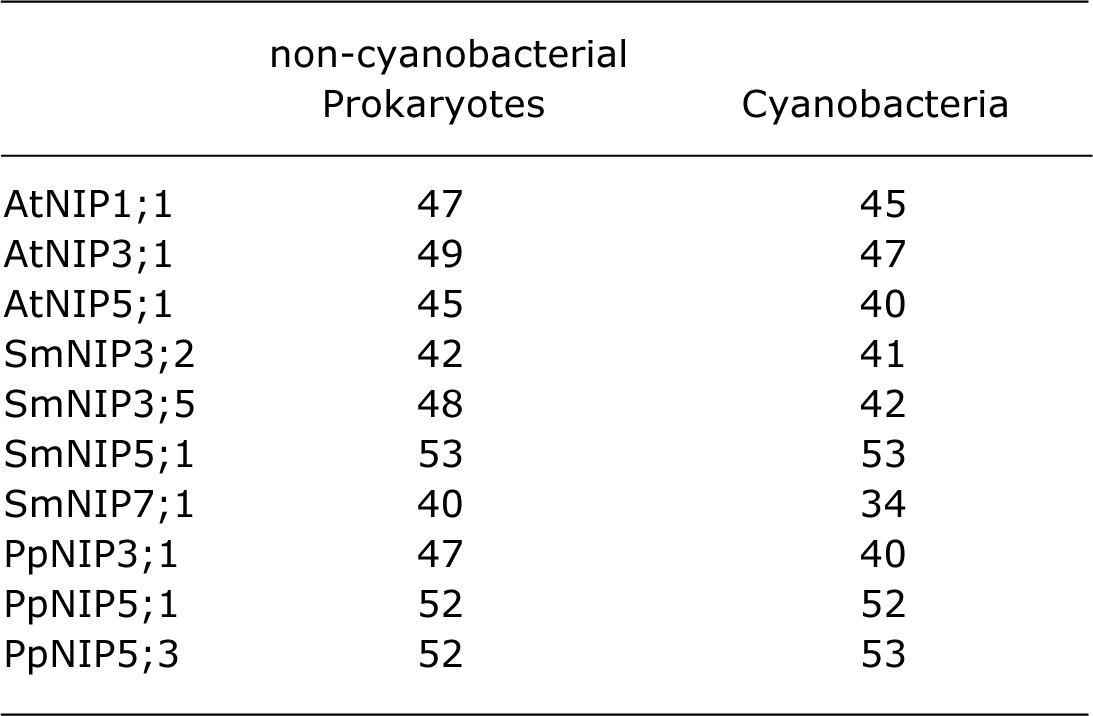
Similarities between NIPs and NIP-likes. Average amino acid identities (%) in three from the five top hits resulting from BLASTP searches against databases (GenBank) of Cyanobacteria, or non-cyanobacterial Prokaryotes, respectively. Queries were NIPs from the angiosperm *Arabidopsis thaliana* (At) [11], the lycopod *Selaginella moellendorffii* (Sm) [13], and the moss *Physcomitrella patens* (Pp) [12]. The three selected proteins were from different genera and coverage of the query protein was at least 70%.

The Archaeplastida comprise the Viridiplantae (green algae and land plants), Rhodophyta (red algae) and Glaucophyta, a small group of unicellular freshwater algae. The superphylum originated from the endosymbiosis of a cyanobacterium and a heterotrophic, eukaryote cell. The cyanobacterium became the plastid, and many of its genes moved to the nucleus of the host cell. About 15% of all plant genes are assumed to be of cyanobacterial descent [36–38], though a much lower estimate of ~2% also has been published [39]. Predictions of cyanobacterial descent are based on sequence comparisons and depend on methodological as well as biological factors, such as the degree of sequence conservation, horizontal gene transfer (between the free-living descendants of the plastid ancestor and other free-living prokaryotes after the plastid arose), and the coverage of the phylum Cyanobacteria [35,38].

In a recent study the coverage of the Cyanobacteria was greatly enhanced by incorporating the genomes of 54 strains [38]. It was predicted that 4339 *Arabidopsis* proteins (13%) are of cyanobacterial descent, including 14 aquaporins: eight out of the nine NIPs, but also half of the ten TIPs and one of the three SIPs. By contrast, none of the 13 *Arabidopsis* PIPs was predicted to stem from cyanobacteria. Similarly, all five NIPs, but also both XIPs and PIP3;1 of the moss *Physcomitrella patens* were predicted to come from cyanobacteria, whereas the remaining aquaporins (7 PIPs, 4 TIPs, 2 SIPs, and 1 GIP) were not [see Supplemental Information]. Indeed, these results may hint at a cyanobacterial origin of NIPs but probably also demonstrate the uncertainty of such predictions.

The transfer of the primordial NIP-gene at the origin of all plants would imply that NIP genes have been present in the forerunners of seed plants, from the last common ancestor of all plants, through their green algal predecessors, to the first land plants. The absence of NIPs in green algae [14], however, seems difficult to reconcile with a vertical transfer of NIP genes in the green lineage since its beginning, and a horizontal transfer after the terrestrialization of plants, which occurred ~700 million years later [40], is still imagined possible [41].

Green algae are divided into two clades, Chlorophyta and Streptophyta [42]. Chlorophytes, the larger group, are unicells, simple multicelluar filaments, or sheet-like and complex corticated thalli of marine, freshwater or terrestrial habitats [42,43]. The streptophyte algae or charophytes, a much smaller group of freshwater algae, are structurally also diverse, from scaly unicellular flagellates through unbranched and branched filaments, to the structurally more complex species of *Chara* (stoneworts) that consist of a main axis with rhizoids and whorls of branchlets [42,44]. All land plants, or embryophytes, (mosses, ferns, horsetails, lycopods, and seed plants) have originated from the streptophyte branch of green algae [44–47]. *Klebsormidium nitens (= K. flaccidum)* is one of the more basal members of the streptophytes. Its thallus consists of simple threads of cells without specialized cells. It is the only charophyte whose genome has been sequenced. Though morphologically quite simple, it has many genes specific for land plants [48]. A BLASTP search at the genome site of *Klebsormidium* revealed it also possesses a very interesting complement of aquaporin genes. It harbors at least 11 aquaporin genes, which can be classified as 5 PIPs, 2 TIPs, 1 SIP, 2 NIPs, and 1 GIP [see Supplemental Information]. Apparently, a NIP gene was already present in the green lineage when the chlorophyte and streptophyte clades diverged, some 800 million years ago [40], but got lost in the chlorophyte branch as no NIPs were found in the chlorophytic algae examined so far [14].

The ancestral gene of plant NIPs probably originated from a cyanobacterium, rather than from a chloroflexian bacterium [30] or some root-dwelling symbiotic bacterium [49]. It is in line with the similarity and abundance of NIP-like genes in cyanobacteria, and it offers a ready explanation for the plant specificity of NIPs. Thus the primordial NIP gene may have entered the green lineage at its very inception by the primary symbiosis of a cyanobacterium and a eukaryotic cell, one of the massive horizontal gene transfers in the history of ife, which is estimated to have occurred ~1500 million years ago [40,50]. A cyanobacterial origin of NIPs is also in good agreement with the estimated dating by Rafael Zardoya and colleagues of ~1200 million years for the origin of the primordial NIP gene, and by inference its transfer to a eukaryotic cell [19].

## Ackowledgement

I wish to thank Jan Jacob Borstlap (Bernotat & Co., Wuppertal, Germany) for the art work.

## References

1. Fortin, M.G., Morrison, N.A., and Verma, D.P.S. (1987). Nodulin-26, a peribacteroid membrane nodulin is expressed independently of the development of the peribacteroid compartment. Nucleic Acids Res. 15, 813–824.

2. Baker, M.E., and Saier Jr., M.H. (1990). A common ancestor for bovine lens fiber major intrinsic protein, soybean nodulin-26 protein, and E. coli glycerol facilitator. Cell 60, 185–186.

3. Preston, G.M., and Agre, P. (1991). Isolation of the cDNA for erythrocyte integral membrane protein of 28 kilodaltons: member of an ancient channel family. Proc. Natl. Acad. Sci. USA 88, 11110–11114.

4. Preston, G.M., Carroll, T.P., Guggino, W.B. and Agre, P. (1992). Appearance of water channels in *Xenopus* oocytes expressing red cell CHIP28 protein. Science 256, 385–387.

5. Kayingo, G., Bill, R.M., Calamita, G., Hohmann, S., and Prior, B.A. (2001). Microbial water channels and glycerol facilitators. Curr. Top. Membr. 51, 335–370.

6. Nehls, U., and Dietz, S. (2014). Fungal aquaporins: cellular functions and ecophysiological perspectives. Appl. Microbiol. Biotechnol. 98, 8835–8851.

7. Campbell, E.M., Ball, A., Hoppler, S., and Bowman, S. (2008) Invertebrate aquaporins: a review. J Comp Physiol B 178, 935–955.

8. Finn, R.N., Chauvigné, F., Hlidberg, J.B., Cutler, C.P., and Cerdà, J. (2014). The lineage-specific evolution of aquaporin gene clusters facilitated tetrapod terrestrial adaptation. PLoS One 9, e113686.

9. Maurel, C., Boursiac, Y., Luu, D.-T., Santoni, V., Shahzad, Z., and Verdoucq V. (2015). Aquaporins in plants. Physiol. Rev. 95, 1321–1358.

10. Laloux, T., Junqueira, B., Maistriaux, L.C., Ahmed, J., Jurkiewicz, A., and Chaumont, F. (2018). Plant and mammal aquaporins: same but different. Int. J. Mol. Sci. 19, 521.

11. Johanson, U., Karlsson, M., Johansson, I., Gustavsson, S., Sjövall, S., Fraysse, L., Weig, A.R., and Kjellbom, P. (2001). The complete set of genes encoding major intrinsic proteins in Arabidopsis provides a framework for a new nomenclature for major intrinsic proteins in plants. Plant Physiol. 126, 1358–1369.

12. Danielson, J.Â.H., and Johanson, U. (2008). Unexpected complexity of the aquaporin gene family in the moss *Physcomitrella patens*. BMC Plant Biol. 8, 45.

13. Anderberg, H.I., Kjellbom, P. and Johanson, U. (2012). Annotation of *Selaginella moellendorffii* major intrinsic proteins and the evolution of the protein family in terrestrial plants. Front. Plant Sci. 3, 33.

14. Anderberg, H.I., Danielson, J.Å.H, and Johanson, U. (2011). Algal MIPs, high diversity and conserved motifs. BMC Evol. Biol. 11, 110.

15. Gustavsson, S., Lebrun, A.-S., Nordén, K., Chaumont, F., and Johanson, U. (2005). A novel plant major intrinsic protein in *Physcomitrella patens* most similar to bacterial glycerol channels. Plant Physiol. 139, 287–295.

16. Wang, W., Haberer, G., Gundlach, H., Gläer, C., Nussbaumer, T., Luo, M.C., Lomsadze, A., Borodovsky, M., Kerstetter, R.A., Shanklin, J., et al. (2014). The *Spirodela polyrhiza* genome reveals insights into its neotenous reduction, fast growth and aquatic lifestyle. Nat. Commun. 5, 3311.

17. Zhang, D.Y., Ali, Z., Wang, C.B., Xu, L., Yi, J.X., Xu, Z.L., Liu, X.Q., He, X.L., Huang, Y.H., Khan, I.A. et al. (2013). Genome-wide sequence characterization and expression analysis of major intrinsic proteins in soybean (*Glycine max* L.). PLoS One 8, e56312.

18. Yuan, D., Li, W., Hua, Y., King, G.J., Xu, F., and Shi, L. (2017). Genome-wide identification and characterization of the aquaporin gene family and transcriptional responses to boron deficiency in *Brassica napus*. Front. Plant Sci. 8, 1336.

19. Zardoya, R., Ding, X., Kitagawa, Y., and Chrispeels, M.J. (2002). Origin of plant glycerol transporters by horizontal gene transfer and functional recruitment. Proc. Natl. Acad. Sci. USA 99, 14893–14896.

20. Bienert, G.P., Schüssler, M.D., and Jahn, T.P. (2007). Metalloids: essential, beneficial or toxic? Major intrinsic proteins sort it out. Trends Biochem. Sci. 33, 20–26.

21. Bhattacharjee, H., Mukhopadhyay, R., Thiyagarajan, S., and Rosen, B.P. (2008). Aquaglyceroporins: ancient channels for metalloids. J. Biol. 7, 33.

22. Pommerrenig, B., Diehn, T.A., and Bienert, G.P. (2015). Metalloido-porins: essentiality of nodulin 26-like intrinsic proteins in metalloid transport. Plant Sci. 238, 212–227.

23. Choi, W.-G., and Roberts, D.M. (2007). *Arabidopsis* NIP2;1, a major intrinsic protein transporter of lactic acid induced by anoxic stress. J. Biol. Chem. 282, 24209–24218.

24. Wang, Y., Li, R., Li, D., Jia, X., Zhou, D., Li, J., Lyi, S.M., Hou, S., Huang, Y., Kochian, L.V., and Liu, J. (2017). NIP1;2 is a plasma membrane-localized transporter mediating aluminum uptake, translocation, and tolerance in *Arabidopsis*. Proc. Natl. Acad. Sci. USA 114, 5047–5052.

25. Grégoire, C., Rémus-Borel, W., Vivancos, J., Labbé, C., Belzile, F., and Bélanger, R.R. (2012). Discovery of a multigene family of aquaporin silicon transporters in the primitive plant *Equisetum arvense*. Plant J. 72, 320–330.

26. Trembath-Reichert, E., Wilson, J.P., McGlynn, S.E. and Fischer, W.W. (2015). Four hundred million years of silica biomineralization in land plants. Proc. Natl. Acad. Sci USA 112, 5449–5454.

27. Zardoya, R. (2005). Phylogeny and evolution of the major intrinsic protein family. Biol. Cell 97, 397–414.

28. Danielson, J.Å.H. and Johanson, U. (2010). Phylogeny of major intrinsic proteins. In MIPs and Their Role in the Exchange of Metalloids, T.P. Jahn and G.P. Bienert, eds. (New York: Landes Bioscience Publishers), pp 19–31.

29. Abascal, F., Irisarri, I., and Zardoya, R. (2014). Diversity and evolution of membrane intrinsic proteins. Biochim. Biophys. Acta 1840, 1468–1481.

30. Finn, R.N. and Cerdà, J. (2015). Evolution and functional diversity of aquaporins. Biol. Bull. 229, 6–23.

31. Zardoya, R., Irisarri, I., and Abascal, F. (2016). Aquaporin discovery in the genomic era. In Aquaporins in Health and Disease. New Molecular Targets for Drug Discovery, G. Soveral, S. Nielsen, and A. Casini, eds. (Boca Raton, Florida: CRC Press), pp 19–31.

32. Calamita, G., Bishai, W.R., Preston, G.M., Guggino, W.B., and Agre, P. (1995). Molecular cloning and characterization of AqpZ, a water channel from *Escherichia coli*. J. Biol. Chem. 270, 29063–29066.

33. Park, J.H. and Saier Jr., M.H. (1996) Phylogenetic characterization of the MIP family of transmembrane channel proteins. J. Membrane Biol. 153, 171–180.

34. Kashiwagi, S., Kanamaru, K., and Mizuno, T. (1995). A *Synechococcus* gene encoding a putative pore-forming intrinsic membrane protein. Biochim. Biophys. Acta 1237, 189–192.

35. Rujan, T., and Martin, W. (2001). How many genes in Arabidopsis come from cyanobacteria? An estimate from 386 protein phylogenies. Trends Genet. 17, 113–120.

36. Martin, W., Rujan, T., Richly, E., Hansen, A., Cornelsen, S., Lins, T., Leister, D., Stoebe, B., Hasegawa, M., and Penny, D. (2002). Evolutionary analysis of *Arabidopsis*, cyanobacterial, and chloroplast genomes reveals plastid phylogeny and thousands of cyanobacterial genes in the nucleus. Proc. Natl. Acad. Sci. USA 99, 12246–12251.

37. Deusch, O., Landan, G., Roettger, M., Gruenheit, N., Kowallik, K.V., Allen, J.F., Martin, W., and Dagan, T. (2008). Genes of cyanobacterial origin in plant nuclear genomes point to a heterocyst-forming plastid ancestor. Mol. Biol. Evol. 25, 748–761.

38. Shih, P.M., Wu, D., Latifi, A., Axen, S.D., Fewer, D.P., Talla, E., Calteau, A., Cai, F., Tandeau de Marsac, N., Rippka, R. et al. (2013). Improving the coverage of the cyanobacterial phylum using diversity-driven genome sequencing. Proc. Natl. Acad. Sci. USA 110, 1053–1058.

39. Makai, S., Li, X., Hussain, J., Cui, C., Wang, Y., Chen, M., Yang, Z., Ma, C., Guo, A.-Y., Zhou, Y. et al. (2015). A census of nuclear cyanobacterial recruits in the plant kingdom. PLoS One 10, e0120527.

40. Yoon, H.S., Hackett, J.D., Ciniglia, C., Pinto, G., and Bhattacharya, D. (2004). A molecular timeline for the origin of photosynthetic eukaryotes. Mol. Biol. Evol. 21, 809–818.

41. Roberts, D.M. and Routray, P. (2017). The nodulin 26 intrinsic protein subfamily. In Plant Aquaporins. From Transport to Signaling. F. Chaumont, and S.D. Tyerman, eds. (Cham: Springer), pp 267–296.

42. Leliaert, F., Smith, D.R., Moreau, H., Herron, M.D., Verbruggen, H., Delwiche, C.F., and De Clerck, O. (2012). Phylogeny and molecular evolution of the green algae. Crit. Rev. Plant Sci. 31, 1–46.

43. Cocquyt, E., Verbruggen, H., Leliaert, F., and De Clerck, O. (2010). Evolution and cytological diversification of the green seaweeds (Ulvophyceae). Mol. Biol. Evol. 27, 2052–2061.

44. McCourt, R.M., Delwiche C.F., and Karol, K.G. (2004). Charophyte algae and land plant origins. Trends Ecol. Evol. 19, 661–666.

45. Becker, B. and Marin, B. (2009). Streptophyte algae and the origin of embryophytes. Ann. Bot. 103, 999–1004.

46. Delwiche, C.F., and Cooper, E.D. (2015). The evolutionary origin of a terrestrial flora. Curr. Biol. 25, R899–R910.

47. de Vries, J., and Archibald, J.M. (2018). Plant evolution: landmarks on the path to terrestrial life. New Phytol. 217, 1428–1434.

48. Hori, K., Maruyama, F., Fujisawa, T., Togashi, T., Yamamoto, N., Seo, M., Sato, S., Yamada, T., Mori, H., Tajima, N., et al. (2014). *Klebsormidium flaccidum* genome reveals primary factors for plant terrestrial adaptation. Nat. Commun. 5, 3978.

49. Ishibashi, K., Morishita, Y., and Tanaka, Y. (2017). The evolutionary aspects of aquaporin family. In Aquaporins, B. Yang, ed. (Dordrecht: Springer), pp 35–50.

50. Parfrey, L.W., Lahr, D.J.G., Knoll, A.H., and Katz, L.A. (2011). Estimating the timing of early eukaryotic diversification with multigene molecular clocks. Proc. Natl. Acad. Sci. USA 108, 13624–13629

